# Revealing the neural representations underlying other-race face perception

**DOI:** 10.1101/2024.09.09.611889

**Authors:** Moaz Shoura, Yong Zhong Liang, Marco Sama, Arijit De, Adrian Nestor

## Abstract

The other-race effect, a disadvantage at recognizing faces of other races than one’s own, has received considerable attention, especially regarding its wide scope and underlying mechanisms. Here, we aim to elucidate its neural and representational basis by relating behavioral performance in East Asian and White individuals to neural decoding and image reconstruction relying on electroencephalography data. Our investigation uncovers a reliable neural counterpart of the other-race effect (i.e., a decoding disadvantage for other-race faces) along with its extended dynamics and prominence across individuals. Further, it retrieves, via neural-based image reconstruction, visual representations underlying other-race face perception and their intrinsic biases. Notably, our data-driven approach reveals that other-race faces are perceived not just as more typical but, also, as younger and more expressive. These findings, pointing to multiple visual biases surrounding the other-race effect, speak to the complexity of its neural mechanisms and its social implications.

## 1. Introduction

The other-race effect (ORE), poorer recognition for faces of a different race than one’s own, has been consistently observed across a diverse array of cultures and races (Hugenberg et al., 2010; Malpass & Kravitz, 1969). Given the scope of its real-world impact, ranging from failures of eye-witness testimony (Wells & Olson, 2001) and identity verification (Sporer et al., 2022) to difficulties with social interactions (McKone et al., 2023), ORE features prominently in the study of visual cognition (for reviews see Ficco et al., 2023; Kubota et al., 2012; Rossion & Michel, 2011). Hence, much has been learned about the relative contribution of perceptual, memory and social factors to its emergence as well as about the neural resources they recruit (Herzmann et al., 2018; Hughes et al., 2019; Natu et al., 2011; Wang et al., 2023; Zhou et al., 2020). Yet, much less is known about the neural representations underlying ORE, their visual content and intrinsic biases.

Accordingly, here, we aim to uncover visual representations involved in other-race (OR) versus same-race (SR) face perception with the aid of neural decoding and image reconstruction applied to electroencephalography (EEG) data (Nemrodov et al., 2018; Nestor et al., 2020). Specifically, we aim to recover and assess the representational structure and the visual content responsible for ORE in East Asian and White participants. Further, we examine the prominence of ORE in neural face processing and its dynamics.

A seminal framework in the study of ORE is that of a psychological face space, a multidimensional construct in which faces are represented as points and their pairwise distances correspond to their perceived similarity (Valentine, 1991; Valentine et al., 2016). In this framework, OR faces are often described as more densely clustered relative to SR faces (Byatt & Rhodes, 2004; Papesh & Goldinger, 2010). Such clustering aims to account for the diminished discriminability of OR faces in experimental settings as well as for the real-world phenomenology of visual experience (e.g., as long described by Feingold: “to the uninitiated American, all Asiatics look alike, while to the Asiatic all white men look alike”; Feingold, 1914). The representational homogeneity of OR faces also finds ground at the neural level, where it elicits repetition suppression, as reported across multiple neuroimaging modalities (Hughes et al., 2019; Zhou et al., 2020). Specifically, viewing pairs of different-identity OR faces may elicit a reduction in neural response comparable to that induced by same-identity face pairs, suggesting that OR faces ‘look alike’ even to the relevant neural population (Reggev et al., 2020; Vizioli et al., 2010).

The density of OR clusters highlight limitations in facial identity information for OR representations and the tendency to reduce them to a prototypical face. However, the optimization of face space dimensions for SR representations (Valentine, 1991) also suggests that OR faces are encoded by suboptimal visual features corresponding to these dimensions (Dahl et al., 2014; O’Toole et al., 1991). Accordingly, a search for ORE-relevant features has led to extensive debates regarding the mis/use of facial shape and surface information (Balas et al., 2011; Balas & Nelson, 2010; Michel et al., 2013; Zhou et al., 2021), and the differential reliance on less holistic processing/features (Harrison et al., 2014; Michel et al., 2006; Tanaka et al., 2004; M. Zhao et al., 2014). Yet, the misrepresentation of OR faces, beyond a reduction in identity information, remains to be elucidated. The present work addresses this challenge by investigating and leveraging the neural counterpart of ORE.

Neural sensitivity to ORE has previously been captured by event-related potentials (ERPs; Tüttenberg & Wiese, 2023), such as enhanced amplitude of the N170 component for OR faces (Senholzi & Ito, 2013; Walker et al., 2008; Wiese & Schweinberger, 2018) – but see (Caldara et al., 2004; Tanaka & Pierce, 2009). Further, functional magnetic resonance imaging (fMRI) studies have reported higher activity in the fusiform face area (FFA) for SR compared to OR faces (Feng et al., 2011; Golby et al., 2001), presumably reflecting the processing of more detailed information for the former in this region. Relevantly here, multivariate fMRI analyses have revealed different spatial activation patterns in the adult ventral cortex for OR versus SR faces (Contreras et al., 2013; Natu et al., 2011; Wang et al., 2023) and have suggested different dynamics (Zhou et al., 2020).

Building on the results above, the present work seeks to address three related aims: (1) relating neural decoding to individual estimates of ORE; (2) characterizing its temporal dynamics, and (3) retrieving and assessing the neural representations underlying ORE. Crucial to our investigation is its reliance on a data-driven approach, which allows uncovering new perceptual biases in OR representations. To anticipate, our results show that: (1) ORE can be reliably related to differences in the neural decoding of OR versus SR faces; (2) the neural dynamics of ORE exhibit an extensive time course, and (3) OR face representations exhibit a typicality bias as well as age and expressiveness biases.

## 2. Materials and methods

### 2.1 Participants

A total of 43 adult (age range: 18-30 years; 28 females) volunteered for the EEG experiment. After excluding three participants (2 East Asian, 1 White) due to technical difficulties with the EEG recordings, 20 identified themselves as East Asian and 20 as White. All participants were right-handed, with normal or corrected-to-normal vision, and no history of neurocognitive disorders.

Prior work (Estudillo, 2021; McKone et al., 2012) on ORE in East Asian and White participants has estimated a medium effect size in both populations (Cohen’s *d* ≥ .62). A power analysis (JASP 0.17.1; jasp-stats.org) for an effect size of .60 (80% power; 5% Type I error rate) indicated that a sample size of 19 participants per group is needed. Thus, our final sample size should allow detecting ORE reliably.

Further, to validate behaviorally our EEG-based reconstruction results, we recruited online (Prolific; www.prolific.co) 50 additional adults (18-35 years; 23 female) – for convenience, referred to below as *validators*. Two were removed because of face recognition scores below the normative range (see below) and two due to failing reliability checks (i.e., negative correlation with themselves across repeated trials). Of the remaining validators, 23 were East Asian and 23 White.

All participants/validators provided informed consent and received monetary compensation. All procedures were approved by the Research Ethics Board at the University of Toronto.

### 2.2 Stimuli

For the EEG experiment we selected 30 images of East Asian males and 30 of White males of similar age, with frontal view and neutral expression, from the Chicago Face Database (Ma et al., 2015). Images were standardized by: (1) aligning the position of the eyes and nose; (2) cropping to display only internal facial features; (3) normalizing with the same mean/contrast values in each CIEL*a*b* channel; and (4) resizing to 98×75 pixels. Resulting stimuli subtended a visual angle of 4°×2.6° from a distance of 80 cm.

### 2.3 Experimental procedures

First, participants completed two versions of the Cambridge Face Memory Test (CFMT) with Chinese (McKone et al., 2012) and White (Duchaine & Nakayama, 2006) face stimuli. This served to assess their face processing proficiency relative to the normative range (i.e., mean ± 2SD, as determined by prior work; Bowles et al., 2009; McKone et al., 2012) and, also, to behaviorally quantify ORE. Both tests have excellent psychometric properties (e.g., Cronbach’s α of .89-.90; McKone et al., 2012; Wilmer et al., 2010) and can be used to estimate individual ORE via the traditional subtraction method (i.e., CFMT score for SR minus that for OR faces; Estudillo, 2021; Wan et al., 2015).

EEG testing was conducted across two 2.5-hour sessions on different days. Participants performed a go/no-go task by pressing a key upon the repetition of the same image on two consecutive trials. Data collection was divided across 32 blocks spanning the two sessions. Within any block, each stimulus was displayed four times, and a random 10% of trials featured a repetition (for a total of 264 trials/block). Stimuli were presented in pseudorandom order (e.g., to avoid repeated “go” trials). Each stimulus was displayed for 300 ms, followed by a variable 600-700 ms inter-stimulus interval during which a center fixation cross replaced the stimulus. Each session began with a practice block. Stimulus presentation and data collection relied on MATLAB and Psychtoolbox 3.0.8 (Brainard, 1997; Pelli, 1997).

### 2.4 Analyses

#### 2.4.1 Behavioral data analysis

Own-race CFMT scores were inspected to confirm that participants exhibit normal face recognition abilities. A two-way ANOVA (2 CFMT versions: East Asian, White; 2 participant groups: East Asian, White) was followed by within-group comparisons (paired *t*-tests) to confirm and compare the size of ORE.

#### 2.4.2 EEG acquisition and preprocessing

EEG signals were recorded with a 64-channel Biosemi ActiveTwo system (Biosemi B.V.). Electrodes were configured according to the international 10/20 system. Offsets were maintained under a threshold of 40 mV. All data were digitally filtered offline (zero-phase 24 dB/octave Butterworth filter) with a bandpass of 0.1–40 Hz and segmented into epochs extending from −100 to 900 ms. Epochs underwent direct current (DC) removal, linear detrending and baseline correction. Noisy electrodes were interpolated if necessary (no more than two electrodes per participant) and epochs were re-referenced to the average reference. Preprocessing relied on Letswave 6 (Mouraux & Iannetti, 2008) and Infomax ICA (Delorme et al., 2007) for artifact removal.

Relatively few trials contained false alarms as participants performed the task at ceiling (mean d’ = 2.73). “Go” trials and those containing artifacts or false alarms were excluded, resulting in an average of 98.1% remaining trials across participants for further analysis.

#### 2.4.2 Pattern classification analyses

We selected 12 electrodes positioned over occipitotemporal (OT) regions (P5, P7, P9, PO3, PO7, O1 on the left, and P6, P8, P10, PO4, PO8, O2 on the right) based on their value in decoding facial information (e.g., as previously determined by multivariate feature/electrode selection; Nemrodov et al., 2020) – for univariate results, see supplementary material.

With respect to the temporal information included in the analysis, two different approaches were considered. First, for temporally-cumulative analyses (Nemrodov et al., 2018; Roberts et al., 2019), we selected a large 50-650 ms time window and, then, concatenated data across electrodes resulting in 3684-feature patterns (i.e., 12 OT electrodes by 307 time points).

The length of this windows aimed to capture both early and late stages of face processing (Ghuman et al., 2014; Nemrodov et al., 2016; Roberts et al., 2019; Vida et al., 2017). Repetitions of all unique stimuli within a given block, timepoint, and OT channel were averaged and z-scored. This yielded a total of 32 observations per stimulus for each participant.

Second, for time-resolved analyses (Dobs et al., 2019; Graumann et al., 2022; Nemrodov et al., 2016), we considered smaller ∼10 ms windows (i.e., five ∼1.95 ms adjacent time bins). Decoding performance was estimated at each position, for a total of 508 intervals, by sliding this window one bin at a time between −100 ms and 700 ms.

Pairwise classification was computed for each participant and stimulus pair by linear SVM (c = 1) with leave-one-block-out cross-validation (across 32 blocks). Classification accuracy for all pairs of facial identities was separately assessed for: (1) SR faces (e.g., White faces for White participants), (2) OR faces (e.g., White faces for East Asian participants), and (3) cross-race faces. Chance-level accuracy was derived by randomly shuffling stimulus labels 1000 times and recomputing classification estimates. Average accuracy across participants was compared to permutation-based chance both for temporally-cumulative analyses (one-sample two-tailed *t*-tests, Bonferroni-corrected across comparisons) and time-resolved analyses (Wilcoxon signed-rank test, FDR-corrected across time points).

#### 2.4.3 EEG-based face space and participant space

To visualize the representational space underlying race perception, we derived an EEG-based face space construct for each group of participants. To this aim, accuracy estimates derived from temporally-cumulative analyses for all pairs of facial identities were averaged across participants in each group and organized into a similarity matrix. Then, metric multidimensional scaling (MDS) was applied to each matrix to derive a face space construct for each participant group (i.e., 40-dimension spaces accounting for at least 90% variance). We anticipated that faces in this space would be clearly separated by race and, also, that OR faces would cluster more tightly than SR ones. To this aim, we computed and compared average pairwise Euclidean distances between each pair of SR faces, each pair of OR faces and each pair of cross-race faces separately for each participant group and corresponding space.

A complementary analysis sought to estimate a participant space and account for its structure. To this aim, principal component analysis (PCA) was applied to EEG-based decoding accuracy vectors across all participants. The first two components of the resulting space were subsequently probed for their relationship with behavioral metrics. Specifically, we assessed the contribution of participant race, facial recognition ability (as captured by SR CFMT scores) and ORE (captured by the difference between SR and OR CFMT scores; Estudillo, 2021; Wan et al., 2015) to this space. To this end, behaviorally derived scores were correlated with PC scores for each dimension.

#### 2.4.4 Image reconstruction procedure

Our reconstruction approach broadly follows earlier efforts (Nemrodov et al., 2018; Roberts et al., 2019; for review see Nestor et al., 2020), which leverage the face space framework to extract visual features and recover neural representations by harnessing the rich spatiotemporal information of EEG patterns (see Fig. 1). Briefly, the procedure involves the following steps: (1) pairwise decoding accuracies for all faces of the same race, except the reconstruction target, were placed into a similarity matrix and averaged across participants, separately for each group; (2) a 20-dimensional face space was derived by applying metric MDS to this similarity matrix; (3) dimension-specific features were computed as classification images through a procedure akin to reverse correlation for all dimensions; (4) feature significance was estimated for each dimension, for each pixel and color channel, via a permutation-based test (i.e., by shuffling stimuli randomly across face space positions 1000 times and recomputing classification images); (5) features that did not contribute any significant pixelwise information in any color channel were eliminated; (6) a target face was projected into the current face space based on its similarity to faces in this space; (7) a reconstruction of the target was achieved by linearly combining significant features, weighted by the target’s coordinates in face space.

**Fig. 1.**
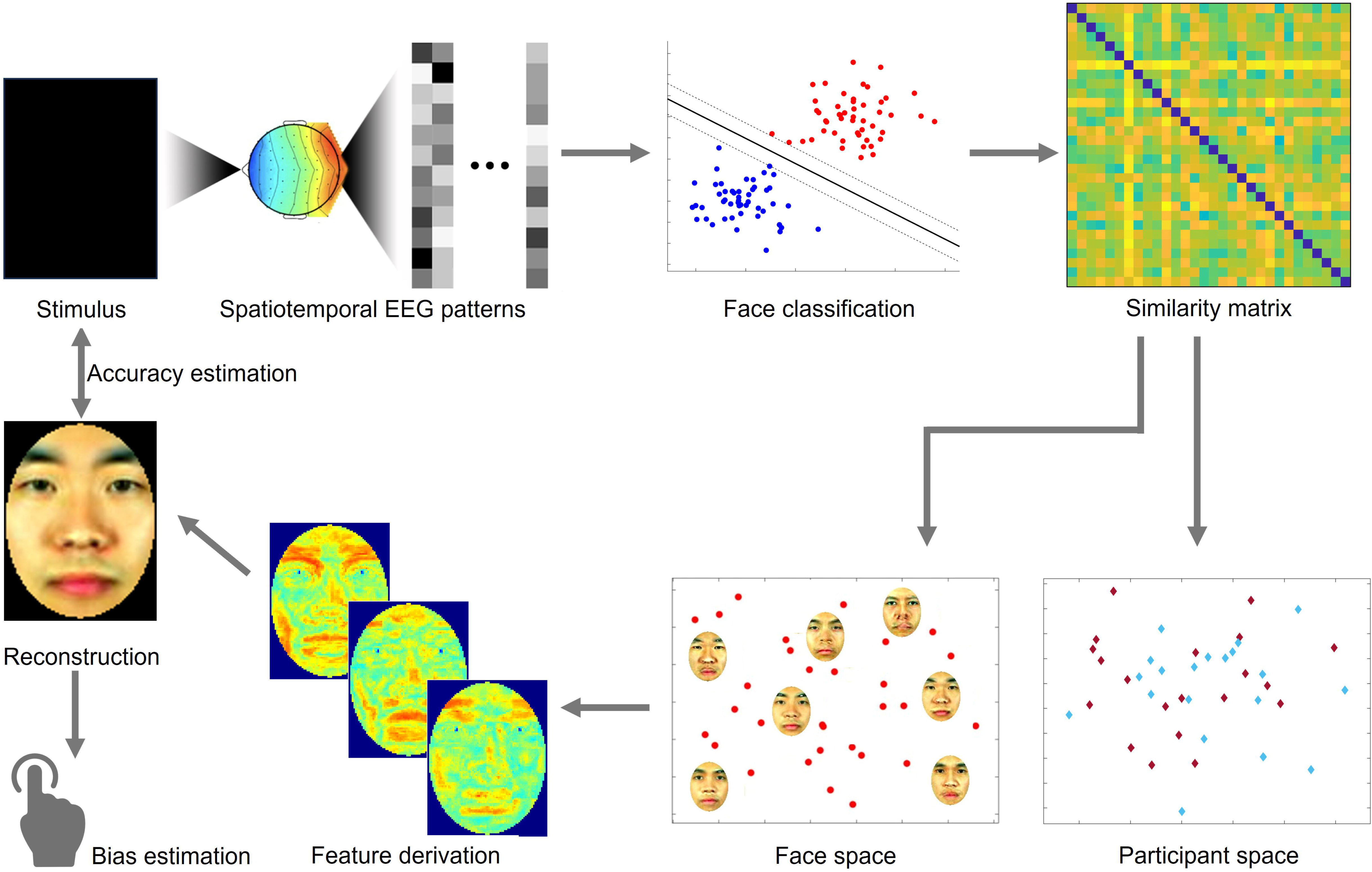
Schematic illustration of data analysis and image reconstruction. Spatiotemporal electroencephalography patterns (EEG) elicited by viewing own- and other-race face stimuli (top left) are linearly decoded and converted into a representational similarity matrix (top right). Pairwise similarity ratings are then converted into EEG-based face space and participant space constructs (bottom right), through multidimensional scaling and principal component analysis, respectively. Facial features are derived directly from the structure of face space and combined into an image reconstruction aiming to recover the neural representation of the corresponding stimulus. The reconstruction is assessed in terms of image accuracy, with respect to its corresponding stimulus, and in terms of potential biases by human observers (bottom left). Stimuli that were taken from the Chicago Face Dataset were removed from the figure due to copyright restrictions. Remaining face images were all artificially generated.

The approach above was applied separately for each face (i.e., by treating each face as a reconstruction target) and relied on a face space derived from the similarity of all other faces sharing the same race with the target (to avoid dependency). By considering decoding patterns for SR and OR faces, this approach aimed to recover both types of visual representations separately in East Asian and White participants.

#### 2.4.5 Image reconstruction evaluation

The accuracy of the image reconstruction for any particular face was assessed by determining the proportion of instances in which a reconstructed image was closer to its target stimulus than to any other face of the same race via a pixelwise L2 distance. To be clear, 100% reconstruction does not indicate a perfect replication of the corresponding stimulus, but only that the reconstruction was more similar to this stimulus than to any other stimulus of the same race. Accuracy was compared against chance (i.e., a one-sample test against 50%) and against each other for reconstructions of different stimulus races using a bootstrap test (10,000 iterations).

To rule out the possibility that any differences between OR and SR reconstructions simply reflect differences in image quality (e.g., due to image blur, pixel noise, spatial distortions) we computed estimates of each reconstructed image via two complementary metrics. Specifically, we appealed to a common reference-based metric, the structural similarity index (SSIM; Wang et al., 2004), as well as to a reference-free metric that approximates perceptual judgements, the blind/referenceless image spatial quality evaluator (BRISQUE; Mittal et al., 2012). OR and SR estimates were then compared to each other using a Wilcoxon signed-rank test separately for each metric and stimulus race.

Last, to assess local, low-level pictorial differences of SR versus OR face reconstructions, we subtracted corresponding images generated from the two groups (i.e., a reconstruction of a given face based on White participants data from a reconstruction of the same facial identity based on East Asian participants data) and assessed their pixelwise significance with a permutation-based test. Specifically, we randomly shuffled the labels of the participants across groups (i.e., East Asian, White), recomputed average similarity matrices, reconstructed face images for each group and subtracted corresponding images generated from the two groups. The initial image differences were then compared with their permutation-based counterparts to identify pixels yielding values different from chance (two-tailed pixelwise permutation test, 1000 permutations; FDR-corrected across pixels, separately for each color channel). This analysis was conducted for every facial identity as well as for the average of all facial identities of the same race.

#### 2.4.6 Behavioral evaluation of reconstruction results

A different group of participants (*validators*) evaluated and compared reconstruction results. First, validators completed the two versions of CFMT, with East Asian (McKone et al., 2012) and White face stimuli (Duchaine & Nakayama, 2006), to assess face processing abilities and ORE.

Then, they viewed 120 image reconstructions (i.e., 30 East Asian faces, reconstructed twice, from each group of EEG participants, and 30 White faces, also reconstructed twice). Since these stimuli are reconstructions of percepts elicited by stimuli in our EEG experiment, their appearance is standardized the same manner (e.g., with respect to size).

On each trial, validators viewed pairs of reconstructions of the same face identity (i.e., derived from East Asian versus White participant data) and selected the face which appeared younger, more expressive, or more typical of its race. Faces were displayed until a response was recorded, with self-paced breaks between blocks to minimize fatigue. Each pair of reconstructions was presented twice, by swapping the left/right position of the reconstructions on the screen. The experiment comprised 6 blocks (3 facial attributes × 2 stimulus races) of 60 trials (30 reconstructions of a given race presented twice). Block and trial order were randomized. Stimulus presentation and data collection relied on PsychoPy (Peirce et al., 2019). Validators completed testing during a 45-minute online session.

After data collection, selection rates were averaged across trials separately by each stimulus race and facial trait for each validator. Selection rates were coded so that values above 50% indicate that reconstructions from OR participants (i.e., White face images reconstructed from East Asian participants, or vice versa) appear younger, more expressive, or more typical of their race. Conversely, a score below 50% indicate that validators judge reconstructions from their own racial group in this manner. Selection scores were then compared to chance (one-sample *t*-tests across validators against 50%, Bonferroni-corrected) and with each other (three-way mixed ANOVA; two validator groups: East Asian and White, two stimulus races, and three facial traits: age, expressiveness, typicality). Last, trait-specific selection rates, averaged across validators, were correlated with each other across all facial identities of the same race (e.g., age and expressiveness rates for East Asian faces) to examine whether judgments capture same/different underlying bias(es) across traits.

## 3. Results

### 3.1 Behavioral performance

Face recognition abilities were assessed using the CFMT with Chinese (McKone et al., 2012) and White (Duchaine & Nakayama, 2006) face stimuli across all participants. An analysis of recognition performance (two-way ANOVA; 2 test versions: Chinese, White × 2 participant groups: East Asian, White) revealed an effect of test version (*F*(1,38) = 10.42, *p* = .003, 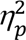 = .22) and participant race (*F*(1,38) = 10.90, *p* = .002, 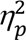 = .22), along with an interaction (*F*(1,38) = 38.1, *p* < .001, 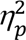 =.50). Post hoc tests revealed that, as expected, White participants outperformed East Asian participants on the CFMT-White (*t*(38) = 5.43, *p* < .001, *d* = 1.72); however, the two participant groups performed comparably on the CFMT-Chinese (*t*(38) = .66, *p* > .999). The latter result may be due to participant background, as ORE tends to be diminished in cities with a diverse multicultural population (Zhou et al., 2022). In our study, a majority (60%) of East Asian participants were international students from a Han Chinese background; in contrast, all White participants were locals from a highly diverse, multicultural city (Toronto, Ontario), whose population is likely to exhibit diminished ORE (Zhou et al., 2022).

More importantly, ORE was confirmed by comparing SR versus OR CFMT scores both for East Asian (two-tailed paired *t*-test, *t*(19) = 5.78, *p* < .001, *d* = 1.29) and White participants (*t*(19) = 2.53, *p* = .02, *d* = .57) - see Fig. 2A. Overall, these results confirm the presence of ORE in both groups, provide evidence for the robustness of our behavioral measures and motivate our investigation into their relationship with neural-based effects below.

**Fig. 2.**
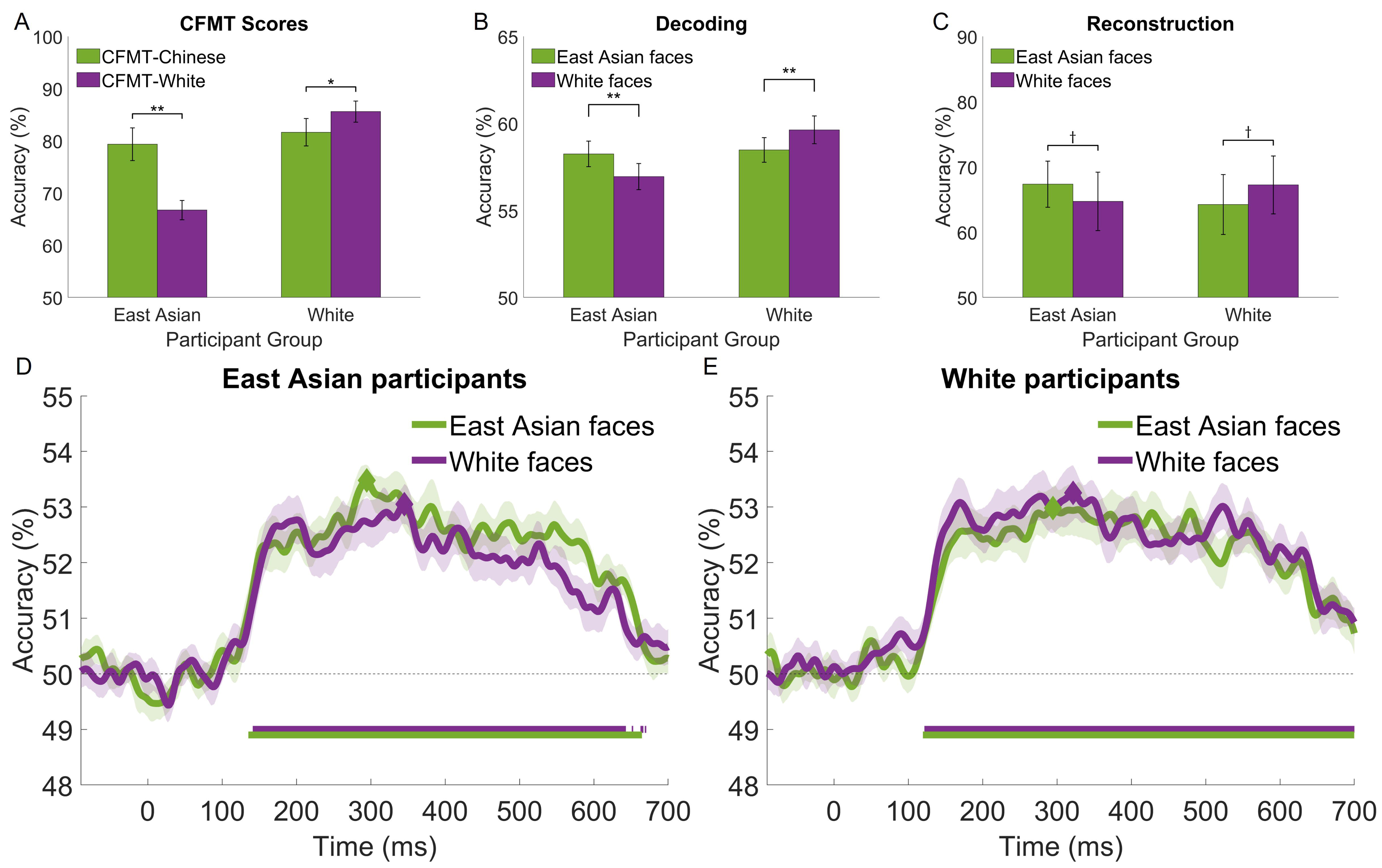
Performance associated with own- and other-race face perception for East Asian and White participants. (A) face recognition accuracy and other-race effects estimated with CFMT-Chinese and CFMT-White; (B) temporally-cumulative pairwise stimulus decoding by stimulus race (based on an 50-650ms interval × 12 electrode patterns); (C) accuracy of EEG-based image reconstructions; (D, E) time-resolved face decoding (based on a 10ms sliding window × 12 electrode patterns) for East Asian and White participants, respectively. Participants exhibit a systematic own-race advantage, which reaches significance (two-tailed *t*-tests across stimulus race) for (A) and (B). The time course of face decoding shows extensive intervals of above-chance decoding (Wilcoxon signed-rank test against permutation-based chance; FDR-corrected across time points, *q* < .05; horizontal bars at the bottom of each plot) and higher estimates of own-race face decoding (though not significant after FRD correction). Diamonds mark the peaks of decoding accuracy in (D) and (E). Error bars for (A), (B), (C) and shaded areas for (D), (E) indicate ± 1 SE (^†^ p<.10, \**p* < .05, ** *p*<. 01).

#### Neural decoding of SR and OR faces

Pairwise face decoding relying on temporally-cumulative decoding (Nemrodov et al., 2018; Roberts et al., 2019) revealed above-chance classification accuracy within and across stimulus race for both participant groups (one-sample *t*-tests across participants against permutation-based chance, all *p’*s < .001, all *d*s > 2.21) – see Fig. 2B. An assessment of classification accuracy (two-way ANOVA; 2 stimulus races × 2 participant groups) revealed no main effects but a significant interaction (*F*(1,38) = 26.98, *p* < .001, η_p_^2^= .42). Subsequent tests indicated that the decoding performance for East Asian face stimuli was comparable in the two participant groups (*t*(38) = .22, *p* = .83). In contrast, White face stimuli were marginally better decoded by White participants compared to East Asian participants (*t*(38) = 2.55, *p* = .058). We note that these results mirror the pattern of behavioral results described above, which we ascribe to participant background.

Importantly, a comparison of SR versus OR face decoding revealed a significant advantage for the former both for East Asian (*t*(19) = 3.29, *p* = .004, *d* = .74) and White participants (*t*(19) = 4.51, *p* < .001, *d* = 1.0). Arguably, these results provide evidence for a neural-based ORE counterpart.

Further, cross-race face decoding yielded higher decoding than within-race decoding both for East Asian participants (relative to White faces: *t*(19) = 8.41, *p* < .001, *d* = 1.88 and East Asian faces: *t*(19) = 3.58, *p* = .002, *d* = .8) and for White participants (relative to White faces *t*(19) = 5.36, *p* < .001, d = 1.2 and East Asian faces: *t*(19) = 6.85, *p* < .001, d = 1.53).

We note that the present results are based on data from two experimental sessions for all participants. Session-specific results, indicative of potential perceptual learning effects, are detailed in the supplementary material. Also, for completeness, analyses were conducted across all electrodes (instead of only 12 OT electrodes) – see supplementary Fig. S1 and S2.

#### Neural dynamics

To evaluate the temporal profile of face processing, decoding was conducted similarly over successive ∼10-ms intervals (instead of a large window spanning most of the trial). For both participant groups and stimulus races, we note above-chance decoding (Wilcoxon signed-rank test against permutation-based chance; FDR-corrected, *q* < .05) starting around 130 ms, peaking between 300-350 ms and tapering off after 600 ms – see Fig. 2D, 2E for East Asian and White participants, respectively.

Interestingly, for both groups, SR faces supported higher decoding accuracy across most time points after 130 ms, though not significantly after multiple-comparison correction (Wilcoxon signed-rank tests across stimulus race, *q* < .05). This suggests that the SR decoding advantage found by temporally-cumulative analysis is not due to any specific, restricted time window. Rather, it likely relies on aggregating complementary information over an extended interval. Accordingly, our results below capitalize on temporally-cumulative results rather than on temporally-restricted ones (e.g., such as those corresponding to decoding peaks).

#### Neural-based face space

Accuracy estimates of pairwise face decoding, averaged across participants, were converted via multidimensional scaling (MDS) into a face space separately for each group (i.e., 40-dimension spaces accounting for at least 90% data variance).

The results evince a clear separation of faces by race (see Fig. 3A, 3B). Additionally, we note a higher degree of clustering for OR than SR faces consistent with face space theory (Valentine, 1991) – in two dimensions this is more clearly apparent for East Asian participants (Fig. 3A). Coarse estimates of pairwise face distances indicate that SR distances are, on average, 2.79% and 1.44% larger than OR ones for East Asian and White participants, respectively. Similarly, cross-race distances are, on average, 2.36% and 4.33% larger than SR ones for East Asian and White participants.

**Fig. 3.**
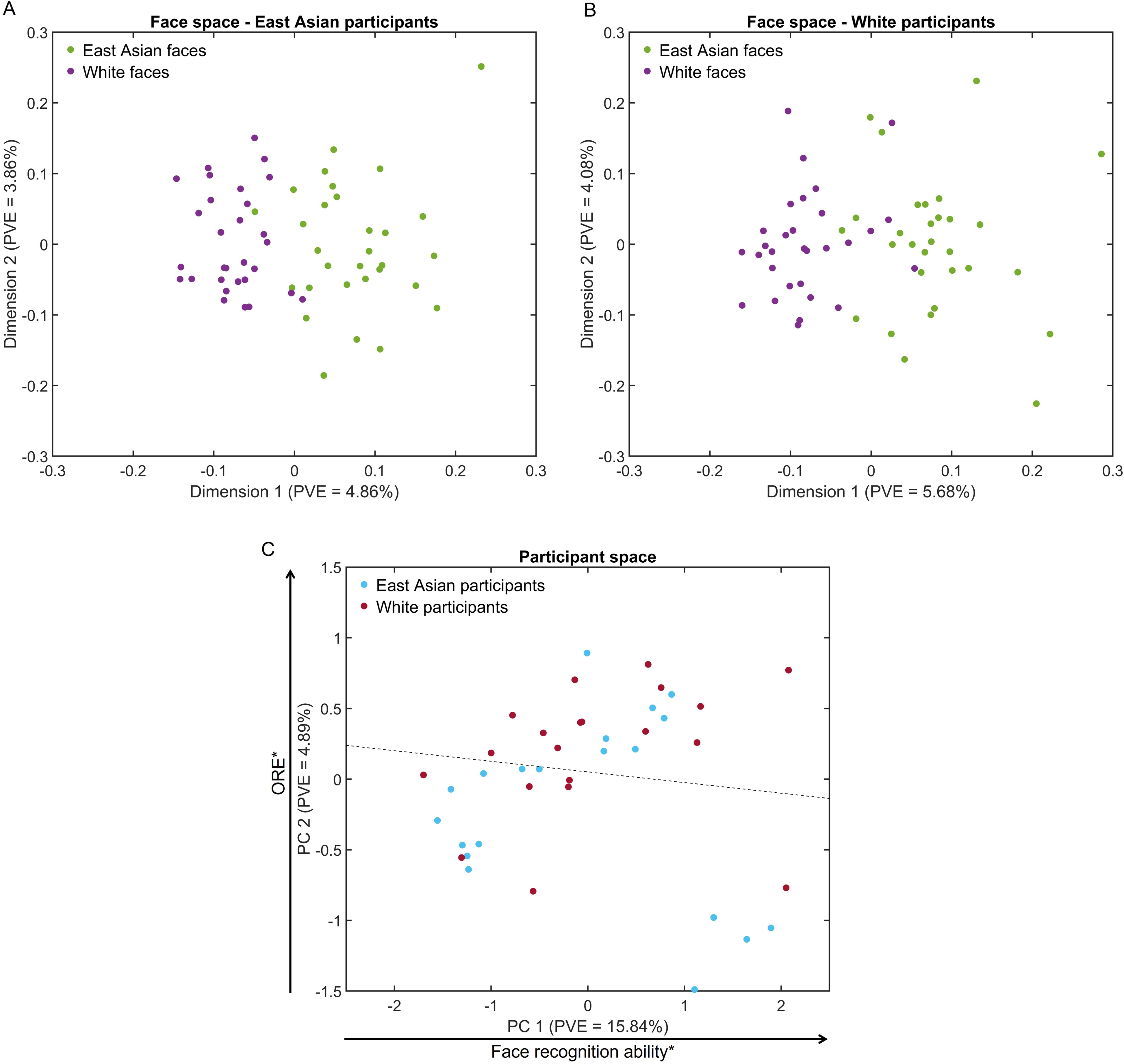
Representational spaces for face stimuli and participants. (A, B) Multidimensional face spaces for East Asian and White participants evince a clear separation of face representations by race and denser clusters for other-race faces (visible in two dimensions for A). (C) A participant space derived from EEG-based similarity vectors evinces some separation of participants by race (the dotted line marks a 65%-accurate hyperplane identified by logistic regression). Notably, participant scores across the first dimension are related to facial recognition ability (Pearson correlation with own-race CFMT across participants) while scores across the second dimension are related to other-race effect estimates (partial correlation with own-race minus other-race CFMT scores while controlling for participant race). PVE – percentage of variance explained (\**p* < .05).

These results reflect the difference in decoding accuracy within and across groups. Specifically, SR faces are better decoded than OR ones leading to larger pairwise distances in face space for the former while cross-race faces are better decoded than within-race faces leading to even larger distances. Interestingly though, we note the prominence of race-based separation (i.e., along the first dimension of face space). These results, based on neural data, complement face space accounts of race informed by behavioral (Byatt & Rhodes, 2004; Papesh & Goldinger, 2010) and computational work (O’Toole et al., 1991; Wang et al., 2023).

To be clear, not all dimensions are likely to be (equally) informative regarding visual face representations and, thus, the estimates above can only serve as a coarse indication of ORE. Hence, a more careful examination of representational content and its sensitivity to ORE has to consider dimension/feature selection (e.g., as implemented by image reconstruction below).

#### Relationship between neural decoding and behavioral performance

Behavioral and neural-based ORE estimates were computed for each participant via subtraction (i.e., SR-OR scores; (Estudillo, 2021; Wan et al., 2015) from CFMT and decoding scores, respectively. These estimates were related to each other across participants from both groups (Pearson correlation, *r*(38) = .68, *p* < .001). To assess whether this correlation reflected within-group differences and not just a categorical difference across participant race, we conducted a partial correlation while controlling for participant race and observed similar results (*r*(38) = .41, *p* = .009). The outcome highlights the relationship between behavioral and neural-based ORE.

Further, an examination of EEG-based participant space, estimated via PCA from pairwise decoding results across participants, revealed some separation between the two participant groups, primarily along the second dimension. To visualize and quantify this separation, participant race was classified in PC space via logistic regression (i.e., across vectors of PCA coefficients corresponding to each participant) and yielded an accuracy of 65% – see Fig. 3C for a separating hyperplane.

To gain more insight into the structure of this space, as reflected by its first two dimensions, we assessed its relationship with estimates of facial recognition: own-race CFMT scores and ORE scores (i.e., SR – OR CFMT scores). We found that PC1 significantly correlated with SR CFMT scores (*r*(38) = .35, *p* = .029), while PC2 evinced a significant correlation with ORE scores (*r*(38) = .37, *p* = .02).

Thus, a neural-based participant space appears to be structured primarily by face recognition ability and ORE. This bolsters the prominence of ORE at the neural level and provides ground for our investigation into the neural representations underlying ORE.

#### Image reconstruction of SR and OR faces

EEG-based image reconstruction (Nemrodov et al., 2018; Nestor et al., 2020) recovered the visual content of SR and OR face representations.

Above-chance reconstruction accuracy was found for both participant groups and stimulus races (two-tailed bootstrap test; all *p’*s < .001). Accuracy for SR faces was higher than for its OR counterpart (Fig. 2C) though the difference was only marginally significant for both East Asian (two-tailed bootstrap test; *p* = .062) and White participants (*p* = .065).

Next, we considered that the lower level of OR reconstruction accuracy may reflect poorer image quality relative to its SR counterpart (e.g., due to loss of high-frequency spatial information or to spatial distortions introduced by the reconstruction method). Accordingly, we evaluated image quality via a common reference-based metric (SSIM) of each reconstruction relative to its corresponding stimulus as well as via a reference-free metric that mimics human perceptual judgements (BRISQUE). A comparison of OR versus SR reconstructions revealed no significant differences by either metric for faces of either race (Wilcoxon signed-rank test; all *p’s* > .40 except for East Asian face reconstructions assessed with BRISQUE, *p* = .072). Hence, we conclude that any differences between OR versus OR reconstructions do not reflect mere image quality differences (e.g., image blur).

Further, to assess local, low-level pictorial differences between SR and OR reconstructions, a pixelwise permutation test was conducted, separately for each facial identity and color channel (two-tailed test, FDR corrected across pixels, *q* < .05). This revealed differences for all channels (Fig. 4A) across multiple facial areas (e.g., around the eyes, eyebrows, nose). The analysis was repeated for image reconstructions averaged across all facial identities of each race (e.g., all Asian face reconstructions based on data from East Asian participants vs. those based on data from White participants) – see Fig. 4B. However, the results are not immediately interpretable in terms of a systematic bias and somewhat less informative for White face reconstructions.

**Fig. 4.**
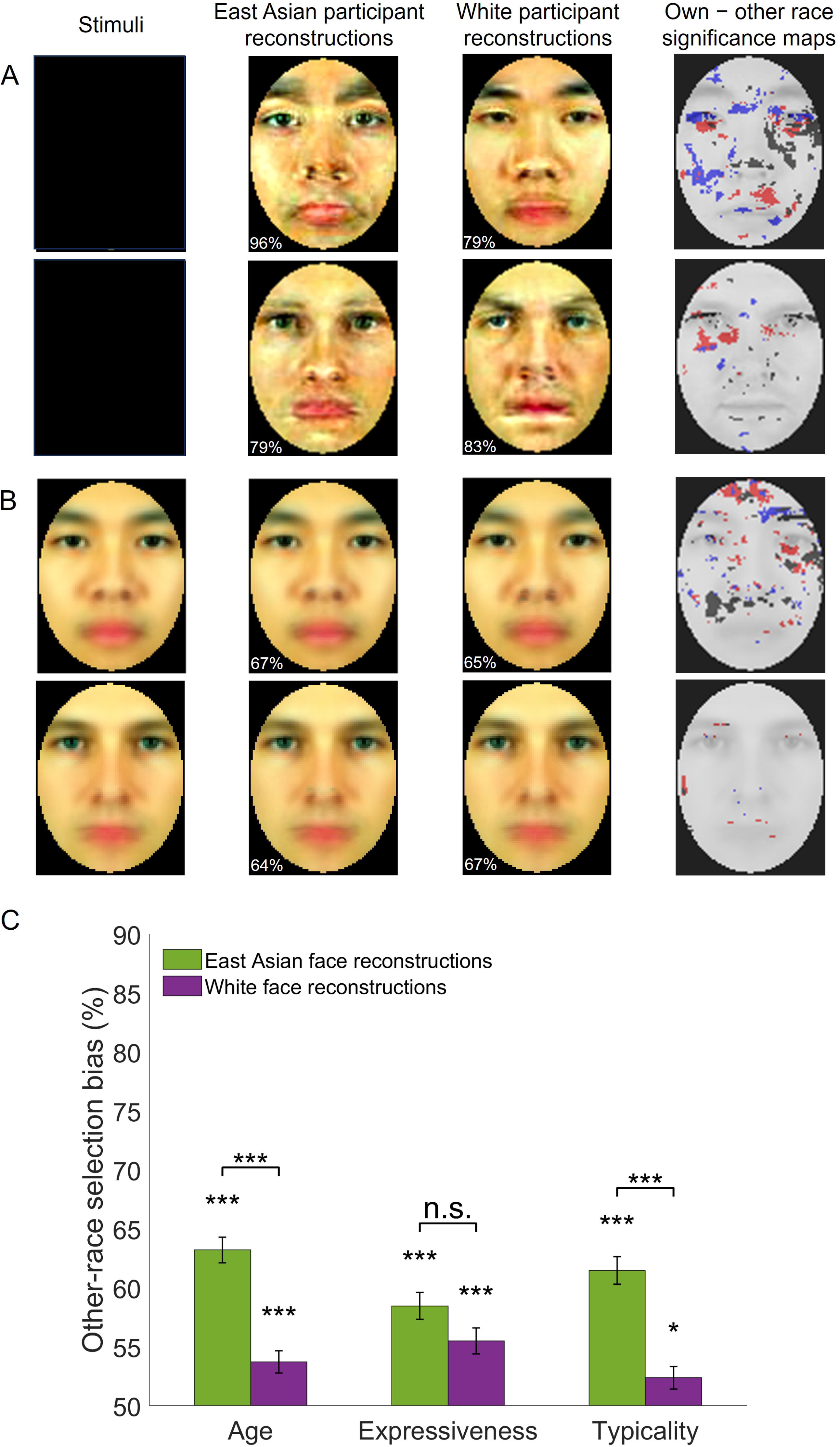
Own-race and other-race facial image reconstructions – examples and assessment. (A) Examples of individual face image reconstructions are shown for East Asian and White participants along with significance maps of the difference between own-versus other-race reconstructions (permutation-based pixelwise test, FDR-corrected, *q* < .05; dark/red/blue pixels mark areas yielding significantly different values in the luminance, red-green and yellow-blue CIEL*a*b* color channels). Numbers in the bottom left corner of each reconstruction indicate accuracy relative to the stimuli in the left column. (B) Averages of all reconstructions of the same race along with corresponding accuracy estimates and significance maps. (C) Other-race reconstructions are judged as younger, more expressive and more typical for their own race relative to own-race reconstructions of the same facial identities. Selection biases are significantly above chance (two-tailed *t*-tests against 50% chance in a 2-alternative forced choice task) and more pronounced for East Asian stimuli (paired *t*-tests for each facial trait). Error bars indicate ± 1 SE (\**p* < .05, \*\*\**p* < .001). Stimuli that were taken from the Chicago Face Dataset were removed from the figure due to copyright restrictions. Remaining face images were all artificially generated.

These results suggest that any divergence between SR versus OR image reconstructions may not be fully captured by low-level visual estimates. Accordingly, next, we asked a separate group of participants to assess the degree of race typicality for each of four sets of reconstructions (i.e., East Asian and White facial image reconstructions recovered from both groups of participants). In addition, we evaluated the possibility that OR visual representations are biased with respect to other facial traits, such as age (Dehon & Brédart, 2001; Mousavi et al., 2018) and expressiveness (Jiang et al., 2023; Yan et al., 2016), and asked validators to also judge these traits.

#### Behavioral evaluation of image reconstructions

For each facial identity, validators viewed and compared its corresponding reconstructions derived from East Asian and White participants. Specifically, on separate trials, they selected the image in a pair that appeared younger, more expressive and more typical for its own race.

Average scores above 50% were noted for all three traits and both stimulus races (one-sample two-tailed *t*-tests, all *p’*s < .001, all *d*s > .58, Bonferroni-corrected). These results indicate that, overall, OR faces are perceived as younger, more expressive, and more typical of their race (Fig. 4C).

An assessment of selection scores (three-way mixed ANOVA; two validator groups: East Asian and White, two stimulus races, three facial traits) found no main effect of group (*F*(1,44) = .46, *p* = .50) or trait (*F*(2,88) = 1.84, *p* = .17) but a main effect of stimulus race (higher scores for East Asian than White faces: *F*(1,44) = 43.23, *p* < .001, η_p_^2^ = .50) as well as an interaction between stimulus race and trait (*F*(2,88) = 7.39, *p* = .001, η_p_^2^ = .14). Subsequent tests indicated that age and typicality scores were significantly more pronounced for East Asian than White face stimuli (*t*(44) = 6.13, *p* < .001, *d* = 1.30 and *t*(44) = 5.87, *p* < .001, *d* = 1.25, respectively) but not expressiveness (*t*(44) = 1.92, *p* = .23).

Next, trait-specific selection rates, averaged across participants, were correlated with each other across all facial identities of the same race (i.e., six correlations for each pair of traits and each stimulus race). All correlations were significant (all *p’s* < .001, except for the correlation between expression and typicality of East Asian stimuli, *p* = .04; Bonferroni-corrected). To assess whether results were entirely driven by typicality, next, we computed partial correlations across age and expressiveness while controlling for typicality. Again, correlations reached significance for both stimulus races (both *p’s* < .001). Last, for completeness, these results were replicated after conducting the analysis separately for each group of validators, East Asian and White (both *p’s* < .001).

Overall, these results indicate that OR faces, regardless of participant race, are perceived as younger, more expressive, and more typical of their race. Last, these biases appear to reflect overlapping but not identical visual sources.

## Discussion

The present work investigates OR face perception with respect to its neural basis, its processing dynamics and its representational visual content. By relating neural decoding and image reconstruction results to behavioral performance in East Asian and White participants this investigation leads to several notable findings.

First, neural decoding, relying on temporally-cumulative occipitotemporal signals, mirrors the ORE evinced by behavioral performance. Specifically, cross-race face decoding is more accurate than within-race decoding, consistent with the separability of neural patterns for OR from SR faces (Natu et al., 2011; Wang et al., 2023). More importantly, within-race face decoding is more accurate for SR than OR faces in both East Asian and White participants. This provides a decoding-based neural counterpart of ORE at the group level (i.e., across participant race), complementing recent results based on repetition suppression (Hughes et al., 2019; Vizioli et al., 2010; Zhou et al., 2020). Furthermore, ORE is also prominent here in neural face processing at the individual level. Specifically, ORE, along with face recognition ability, accounted for the main components of variability in representational similarity across participants. Overall, these results speak to the neural basis of ORE while they also serve as a platform for the rest of our investigation.

Second, the temporal profile of decoding exhibits an extensive window of significance, between around 130 - 600 ms, consistent with prior work on the neural dynamics of facial identity perception (Ambrus et al., 2019; Dobs et al., 2019; Kovács et al., 2023; Nemrodov et al., 2018). Overall, this profile is similar for SR and OR faces as no specific time points, in isolation, yield a significant difference. However, an SR decoding advantage is, at least, numerically apparent during most of the 130-600 ms interval (Fig. 2D, E). This suggests the presence of complementary information over time, which accrues into an overall SR advantage, captured by temporally-cumulative analysis. These results are also consistent with multiple stages of neural processing across an extensive network (Natu et al., 2011; Zhou et al., 2020). While an ORE univariate effect for the N170 ERP component (Balas & Nelson, 2010; Senholzi & Ito, 2013; Walker et al., 2008; Wiese & Schweinberger, 2018) is also apparent (see supplementary material), our decoding results indicate that ORE is unlikely to be solely linked to a restricted temporal window (e.g., only around the N170 component).

Third, neural face representations, recovered through EEG-based image reconstruction, revealed visual differences between SR and OR faces which do not reflect mere differences in image quality. Specifically, image reconstructions assessed by a separate group of validators, revealed that OR faces are represented as more typical for their race. This agrees with the OR compression in face space (Byatt & Rhodes, 2004; Papesh & Goldinger, 2010), which we also find here based on EEG data, as well as the phenomenology of ORE (i.e., “they all look the same”; Ackerman et al., 2006; Feingold, 1914; Laurence et al., 2016). Notably though, we also find largely separate biases in the perception of age and expressiveness, which we address next.

Little is known about OR biases in age perception, with a handful of studies yielding conflicting results (Dehon & Brédart, 2001; Mousavi et al., 2018; L. Zhao & Bentin, 2008). However, an *illusion* of Asian youthfulness is suggested by race-specific differences, relative to White faces, in both skin physiology (e.g., more collagen, thicker dermis) and skeletal structure (Shirakabe et al., 2003; Vashi et al., 2016). Hence, we reasoned that Asian faces may be perceived as younger by White participants, who have less visual experience in discounting the contribution of such factors to the appearance of youth. Our results support this hypothesis but, interestingly, they also show a reciprocal bias, with Asians perceiving White faces as younger. Thus, the present findings speak to a more general OR bias in age estimation and, critically, they reveal the visual representations supporting prior reports of such biases (Dehon & Brédart, 2001; Mousavi et al., 2018).

Regarding expressiveness, recent work has found poorer performance in expression recognition for OR faces (Jiang et al., 2023; Yan et al., 2016), which may be driven by cultural differences in the representation of emotional expressions (Chen et al., 2024; Jack et al., 2012). While our stimuli and corresponding reconstructions display neutral expressions, emotion can be perceived even in neutral faces as a function of facial structure (Neth & Martinez, 2009; Said et al., 2009) and person knowledge (Suess et al., 2015). Hence, we reasoned that biases in expression recognition may also extend to the degree of expressiveness perceived in OR neutral faces. The results bore out this hypothesis and, again, they revealed the neural representations underlying this bias. Further work though will be needed to uncover the precise nature of this bias (e.g., as driven by valence, arousal).

Relating visual biases back to the face space framework, one should consider their underlying source and degree of separability. One possibility is that they all reflect OR clustering in face space. For instance, the appearance of youth may be an outcome of typicality (e.g., by averaging across multiple individuals and reducing the prominence of aging cues). However, we find that, age and expressiveness were correlated even after controlling for typicality. Further, OR biases were less pronounced for White faces despite their denser clustering (and poorer discrimination) compared to East Asian faces. Thus, most likely, OR biases reflect the interplay between multiple properties of face space besides cluster density: the appearance of an OR face prototype (Jaquet et al., 2008; Zhou et al., 2016), the visual features used for representation (e.g., corresponding here to significant dimensions recruited by reconstruction) as well as the topography of the space across subsets of relevant dimensions. Overall, current findings may reflect face space optimization for SR young adult faces (Zhou et al., 2016).

While our examination of potential biases was driven by specific hypotheses, current results may be further queried for other facial trait biases (e.g., attractiveness, competence). Methodologically, this opportunity illustrates the benefit of data-driven approaches aimed at recovering internal representations (Chen et al., 2024; Nestor et al., 2016, 2020; Zhan et al., 2019), including their ability to uncover new perceptual biases. Further, theoretically, it showcases how encoding OR faces in a suboptimal face space (Dahl et al., 2014; O’Toole et al., 1991), crafted for SR recognition, may lead to an array of representational distortions impacting multiple facial traits. In turn, practically, such an array of biases can shed new light on an ORE-induced decrement in the quality of social interactions (McKone et al., 2023), beyond difficulties with person identification.

As a caveat, our present results are likely to reflect automatic perceptual processes associated with brief viewing of facial stimuli, rather than higher-level (e.g., conceptual) processes. While prior work argues for the contribution of memory (Herzmann et al., 2022; Tanaka et al., 2004; Yaros et al., 2019; Zhao et al., 2014) and socio-cognitive factors (Hugenberg et al., 2010; Schwartz et al., 2023) to ORE, our results point to the grounding of ORE in perception (Megreya et al., 2011). However, the main source of ORE may well vary across different groups. For instance, it may be driven by perceptual experience for Asian-White groups, as suggested here, but by social-motivational factors across Black-White groups (Wan et al., 2015). Accordingly, investigating the nature and extent of visual biases across other groups (e.g., Black-White) could be quite informative in this respect. Further, evaluating memory-based representations (Chang et al., 2017; Zhan et al., 2019) and their biases in ORE could help clarify the relevant interplay between perception and memory.

Related to the point above, we also note the reduction in the neural counterpart of ORE across different-day experimental sessions (see supplementary material, Neural decoding and Supplementary Fig. 2), which may be due to perceptual learning (Heron-Delaney et al., 2011; Tanaka & Pierce, 2009; Yovel et al., 2012). The ability of visual experience to reduce visual and social bias, even with limited perceptual training (Lebrecht et al., 2009), is a topic of significant theoretical and practical interest. Hence, investigations methodologically similar to the present one but targeting perceptual learning may offer valuable insights into how visual experience alters the neural representations underlying ORE and reduce bias.

To conclude, the present work integrates measures of behavioral performance, neural decoding and image reconstruction to provide new insights into the representational basis of ORE and its dynamics. Our findings reveal multiple biases in OR face perception with significant theoretical, methodological and practical implications. More generally, they open new avenues for exploring racial biases and their scope in face recognition.

## Author contributions

**Moaz Shoura:** Conceptualization, Methodology, Software, Formal Analysis, Investigation, Data curation, Validation, Visualization, Writing – Original Draft/Review & Editing; **Yong Zhong Liang:** Formal Analysis, Investigation, Data curation, Writing – Original Draft; **Marco Sama**: Formal Analysis, Investigation, Data curation, Writing – Original Draft; **Arijit De**: Investigation, Data curation, Writing – Original Draft; **Adrian Nestor:** Conceptualization, Formal Analysis, Methodology, Validation, Original Draft/Writing-Reviewing and Editing, Resources, Supervision, Funding acquisition.

