## Supplemental information for "Revealing the neural representations underlying other-race face perception"

### Supplementary Materials

#### Univariate ERP results

Extensive research and debate have targeted the sensitivity of the N170 ERP component to the other-race effect (ORE), with multiple studies reporting larger amplitudes for OR than SR faces (Balas & Nelson, 2010; Walker et al., 2008; Wiese & Schweinberger, 2018). However, other studies reported no differences (Caldara et al., 2004; Tanaka & Pierce, 2009) or task-dependent effects, such as larger OR amplitudes when participants attend to facial identity but lower when they attend to race (Senholzi & Ito, 2013).

Accordingly, here, we also assessed the modulation of N170 amplitude and latency by ORE. To this end, average ERP signals were separately computed for 6 left and 6 right occipitotemporal (OT) electrodes (see Methods, Pattern classification analyses), which evince a robust N170 component. The data were further averaged across different face stimuli by race. Three participants (one East Asian, two White) who yielded outlier values for N170 amplitude and/or latency (i.e., beyond ±2.5 SD from the mean of the other participants in their group) were excluded from further analyses. A three-way mixed-design ANOVA (JASP 0.17.1; jasp-stats.org) was conducted across remaining participants separately for amplitude and latency estimates, with participant group (East Asian, White) as a between-subjects factor, and stimulus race (East Asian, White) and lateralization (left, right electrodes) as within-subjects factors.

The analysis of N170 amplitudes revealed a significant main effect of stimulus race (larger amplitude for White faces relative to East Asian ones; $F(1,35) = 6.54$, $p = .015$, $\eta_p^2 = .16$) but no significant effects of participant group, lateralization or interactions (all *p's* > 0.1). However, further examination revealed that the effect of stimulus race was only present for East Asian (t(18) = 2.39, *p*=.03, *d* = .55) but not for White participants (t(17)= 1.07, *p* = .30, *d* = .25). Hence, the N170 component evinces higher amplitude for OR faces in one participant group. While this effect is not significant in our White group, this may be due to the somewhat lower level of ORE found for this group (see Results, Behavioral performance). No main effects or interactions were significant for a similar analysis of latency values (all *p's* > .05). These results are largely consistent with prior work and, also, suggest that participants attend to facial identity rather than to facial race (Senholzi & Ito, 2013), accounting for our ability to discriminate within-race faces via neural decoding.

We note that several other ERP components (e.g., P1, P2, N250) also carry relevance for the study of ORE (for a recent review see Tüttenberg & Wiese, 2023). However, a thorough assessment of their univariate sensitivity to ORE is beyond the main goals of the present work and was not examined further.

**Neural decoding: all-electrode and session-specific results**

Given the possibility that multiple electrode sites across the scalp may be sensitive to ORE (Tüttenberg & Wiese, 2023), in addition to the 12 OT subset considered in our main results, further analyses were conducted using all 64 electrodes. An assessment of classification accuracy (two-way ANOVA; 2 stimulus races × 2 participant groups) revealed no main effects or interaction (all *p's* > 0.05) – see Supplementary Fig. S1. While performance was significantly above chance in all cases (one-sample two-tailed t-tests against permutation-based chance; all *p's* < 0.05), we note a systematic decrement in accuracy (i.e., 1-2.5%) relative to the results based on 12 OT electrodes (Fig. 2B). This likely reflects the cost of increasing the dimensionality of classification patterns which is not offset by the addition of complementary information from other electrodes. More importantly, we found no evidence for a neural-based counterpart of ORE (two-tailed paired t-tests across stimulus race for each participant group; all $p$'s > .10). Thus, our analyses focus on results based on the 12 OT electrode subset.

Next, temporally-cumulative analysis was conducted separately for sessions 1 and 2 to investigate the impact of potential perceptual learning (Heron-Delaney et al., 2011; Hills & Lewis, 2006; Tanaka & Pierce, 2009) across sessions. To this end, classifier training and testing was conducted only on blocks from the same experimental session of each participant. Again, here, performance was significantly above chance in all cases (one-sample two-tailed t-tests against permutation-based chance; all p's < .05) for both sessions – see Supplementary Fig. S2. However, we also note a systematic decrement in accuracy for either session (i.e., 1-3%) relative to the results based on both sessions (Fig. 2B). This likely reflects the smaller training set associated with session-specific decoding. Also, it supports our decision to combine data from two sessions per participant to provide more reliable estimates of decoding for our main set of results.

Interestingly though, for session 1 an SR decoding advantage (two-tailed paired t-test across stimulus race) was present for both East Asian participants ($t(19) = 2.72$, $p = .014$, $d = 0.61$) and White participants ($t(19) = 3.29$, $p = .004$, $d = 0.74$) while this effect was no longer present for session 2 (East Asian participants: $t(19) = .88$, $p = .39$; White participants: $t(19) = -1.57$, $p = .13$). These results may reflect the outcome of perceptual learning for other-race stimuli across the two sessions, leading to a reduction in ORE, at least with respect to the present stimulus set. However, these results need to be interpreted with caution given the more limited number of trials used to estimate session-specific decoding.

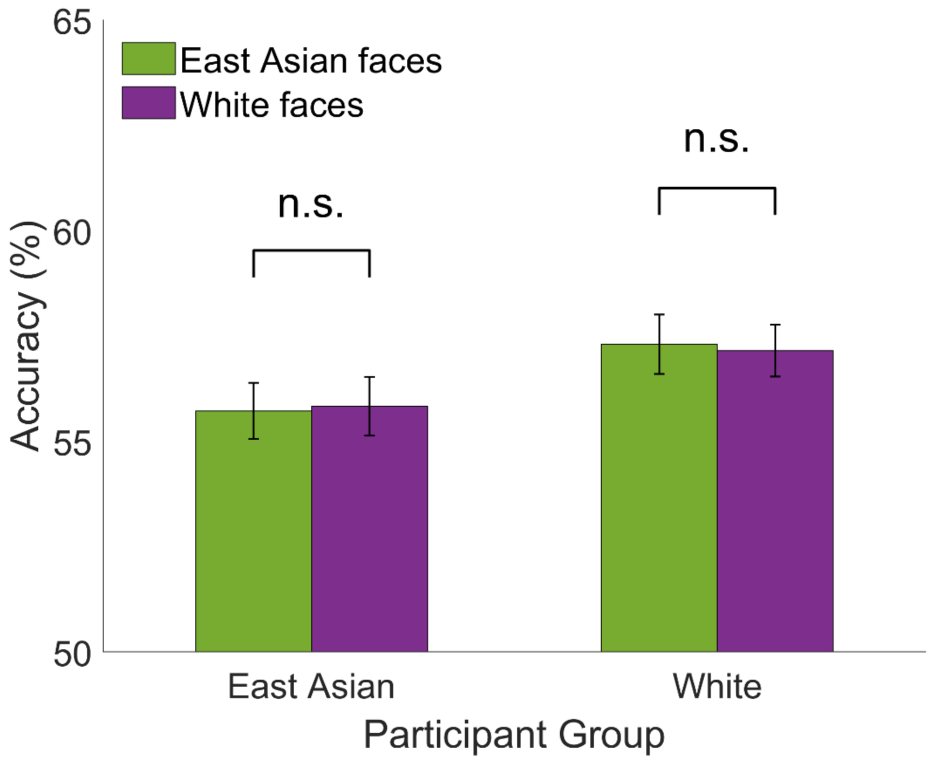


Supplementary Fig. S1. Temporally-cumulative stimulus decoding by stimulus race and participant group. Decoding relies on a 50-650ms interval after stimulus onset and all 64 electrodes. Decoding performance is poorer relative to that based on 12 OT electrodes (Fig. 2B) and does not evince an own-race advantage (two-tailed t-tests across stimulus race for each participant group, both $p$'s $>.10$).

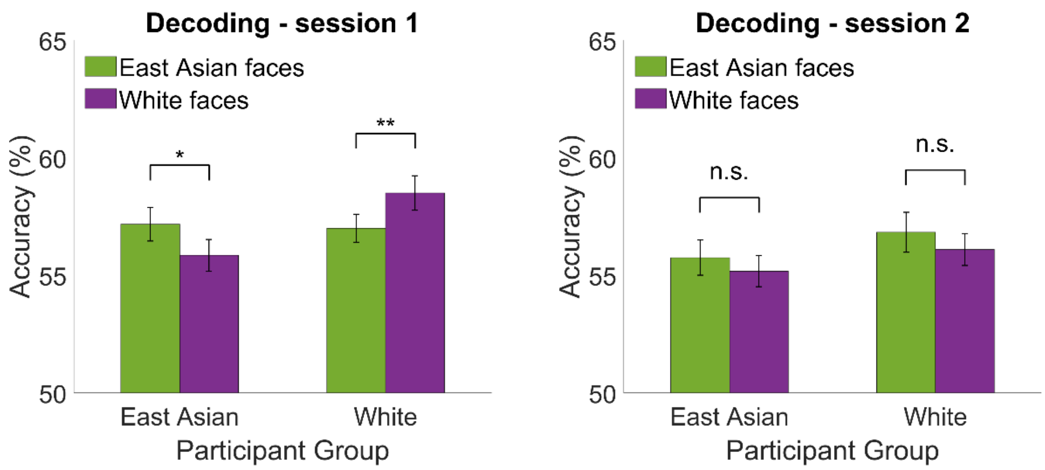


Supplementary Fig. S2. Temporally-cumulative stimulus decoding by stimulus race and participant group, separately for (A) session 1 and (B) session 2. An own-race advantage is present for each participant group only in session 1 (two-tailed t-tests across stimulus race for each participant group (*$p$<.05, **$p$<.01).
